# Multidimensional natal isotopic niches reflect migratory patterns in birds

**DOI:** 10.1101/2021.04.27.441573

**Authors:** A. Franzoi, S. Larsen, P. Franceschi, K.A. Hobson, P. Pedrini, F. Camin, L. Bontempo

## Abstract

Naturally occurring stable isotope ratios in animal tissues allow estimation of species trophic position and ecological niche. Measuring multiple isotopes of migratory species along flyway bottlenecks offers the opportunity to sample multiple populations and species whose tissues carry information at continental scales.

We measured *δ*^2^H, *δ*^18^O, *δ*^13^C, *δ*^15^N in juvenile feathers of 21 bird species captured at a migratory bottleneck in the Italian Alps.

We examined if trends in individual isotopes reflected known migratory strategies and whether dietary (*δ*^13^C-*δ*^15^N) and spatially-explicit breeding origin (*δ*^2^H-*δ*^18^O) niche breadth (NB) differed among long-distance trans-Saharan (TS), short-distance (IP) and irruptive (IR) intra-Palearctic migrants, and whether they correlated with reported populations long-term trends.

In both TS and IP groups, species *δ*^2^H declined with capture date, indicating that northern populations reached the stopover site later in the season, following a Type-I migration strategy. Values of *δ*^2^H indicated that breeding range of TS migrants extended farther north than IP and IR migrants. The breeding season was longer for IP migrants whose *δ*^13^C and *δ*^15^N values declined and increased, respectively, with time of capture.

Average species dietary NB did not differ among migratory groups, but TS migrants displayed wider breeding origin niches, suggesting that long-distant migration is linked to broader ecological niches. Isotope origin NB well reflected species geographic range extent, while dietary NB did not correlate with literature accounts of species’ diet. We found no relationship between species breeding NB and population trends in Europe, implying that conditions in the wintering grounds could be more important for the long-term persistence of species.

We demonstrate that isotopic measurements of passerines migrating through a bottleneck represents a unique opportunity to investigate large-scale life-history phenomena relevant to conservation.

## Introduction

The concept of species’ niche is a fundamental pillar of ecological science (Hutchinson, 1957; Chase and Leibold, 2003; Holt, 2009), but remains challenging to quantify empirically. The challenge is particularly acute in the case of long-distance migratory birds due to the seasonal shifts in habitat use and distribution throughout their life-cycles. Yet, habitat conditions in the breeding, stop-over and wintering grounds profoundly influence individual fitness and demography of migratory populations. Long-distance migrants are showing rapid population declines and appear at greater risk of extinction compared to residents (Heldbjerg and Fox, 2008; Evans et al., 2012; Vickery et al., 2014). Therefore, assessing niche parameters and differences among migratory groups can help in understanding the drivers shaping migration strategies and setting conservation actions.

Migratory pathways of multiple species often overlap at stopover sites, which can represent bottlenecks, or passage points, causing high spatio-temporal concentration of many individuals and species from different migratory systems (Heiss, 2013; Buechley et al., 2018; Cardenas-Ortiz et al., 2020). Therefore, estimating niche parameters of individuals intercepted along migratory flyways represents a valuable and cost-effective approach to include multiple species potentially conveying information from large, continental scale, breeding areas (Yohannes et al., 2007; Hobson et al. 2015; Heim et al. 2018; Cardenas-Ortiz et al., 2020).

Ecological niches of species can be approximated using measurements of naturally occurring stable isotope ratios in their tissues. Such “isotopic niches” can be used to empirically quantify the “realised” niche dimensions of species (Bearhop et al., 2004; Newsome et al., 2007), since stable isotope measurements in animal tissues provides a time-integrated average of many feeding events (Hobson 2011). In particular, stable isotope ratios of carbon (*δ*^13^C) and nitrogen (*δ*^15^N) have been used to represent the “dietary niche” of an organism because they reflect reliance on different primary producers and trophic level, respectively. Although, the dietary niche breadth represented by δ^13^C and δ^15^N reflects the range in isotope values of dietary inputs in any given area (Hoeninghaus and Zueg 2008, Hette-Tronquar 2019), the isotopic niche space produced is a potentially powerful means of deriving information on diets of migratory birds from broad catchment areas (Newsome et al. 2007, Inger and Bearhop 2008).

Similarly, isotopic values of hydrogen (deuterium; δ^2^H) and oxygen (δ^18^O) are linked to isotopic values in environmental waters that can show predictable spatial patterns or “isoscapes” at continental scales (Bowen, 2010). Therefore, tissue δ^2^H and δ^18^O values can be used to represent a spatially-explicit “origin niche” of bird species (e.g. Hobson et al. 2004). Although the meteoric relationship predicts a strong correlation between water δ^2^H and δ^18^O, this strong relationship tends to break down in consumer tissues within foodwebs due to the fact that oxygen in tissues, unlike hydrogen, can be derived from air as well as diet and drinking water. Moreover, metabolic pathways leading to oxygen incorporation in proteins differ from those involving hydrogen (Vander Zanden et al. 2016, Magozzi et al. 2019). The net result is that the two isotopes tend to be poorly correlated in feathers (Hobson and Koehler 2015). As such, values of δ^2^H and δ^18^O could be considered independent in the analyses of feathers and their combined isotopic space would be highly associated with the geographic origins where feathers were grown (Pekarsky et al. 2015).

The use of multiple isotopes in animal tissues has allowed ecologists to estimate trophic niche breadths of different species or individuals in populations (Bearhop et al., 2004; Shipley and Matich, 2020), and the geographic origin of migratory species (Hobson, 1999; Hobson and Wassenaar 2019). These studies have provided insight into trophic relationships in communities (Post, 2002; Abrantes et al., 2014; Wang et al., 2018) as well dietary specialisation (e.g. Bearhop et al. 2004) and migratory pathways of declining bird species (Evans et al., 2012; Steenweg et al., 2017). Due to logistic and economic constraints, most studies measuring both dietary and origin isotope niches have focused on single species or local populations and communities (but see Yohannes et al., 2007; Rader et al., 2017; Ma et al. 2020). However, examining multiple isotope niches at migratory bottlenecks offers the unique opportunity to quantify and compare niche parameters among several bird species that evolved different migratory behaviours.

In this study, we measured multiple isotopes in the juvenile feathers of approximately 700 individuals from 21 species of European passerines captured during the autumn migration at an Alpine migratory bottleneck over the Italian Alps. Metabolically inert tissues such as feathers show an isotopic composition that reflects the environment during their formation. As a consequence, bird feathers grown in the nest represent “remote carriers” of isotopic information from the breeding grounds (Langin et al., 2007), since many species retain juvenile plumage during subsequent migration and overwintering. In particular, we examined whether dietary (*δ*^13^C-*δ*^15^N) and origin (*δ*^2^H-*δ*^18^O) niche breadths (NB) differed among species belonging to the *trans-Saharan* (TSM), *intra-Palearctic* (IPM), and *irruptive intra-Palearctic* (IRM) migratory systems, and the extent to which feather stable isotope values reflected known migratory patterns.

For both TSM and IPM groups, we expected northern populations to reach the autumn Alpine passage later in the season. According to the deuterium precipitation isoscape in the Palearctic (Hobson et al., 2004; Bowen, 2010), this would result in a negative relation between arrival date and *δ*^2^H for each species, as also observed in Neotropical species adopting a Type-I migration strategy whereby southern populations precede northern ones or as “chain migrants” (Cardenas-Ortiz et al., 2020).

Since IPM species display a longer breeding season (2-3 clutches) compared to TSM (1-2 clutches), we also compared seasonal variation in *δ*^13^C and *δ*^15^N for each species to examine seasonal changes in the availability of main food sources (e.g. larvae to adult insects).

Long-distance migration is thought to have evolved primarily in species able to utilise multiple habitat types (Levey and Stiles, 1992; Cresswell, 2014; Reif et al., 2016), which are encountered *en route* and in wintering areas. More generally, if migratory connectivity occurs at large scales, it could favour the selection for generalist traits. As such, species migration distance is expected to be directly related to their niche breadth. Although intuitively appealing, this hypothesis has received limited empirical support so far (Cresswell, 2014; Laube et al., 2015; Ponti et al., 2020). Here, we specifically examine how migratory distance was related to the origin niche breadth (*δ*^2^H-*δ*^18^O). We test the hypothesis that long-distance migrants such as TSM species, display broader niches than IPM and IRM species.

Because the relation between isotope niche measures and other estimates of species niche are still poorly understood (Shipley and Matich, 2020), we also examined whether our measures of dietary and origin isotope niches matched diet and climatic niche breadth derived from the literature and species range maps (Reif et al., 2016). While we expected estimates of spatially-explicit isotope origin niche to accord with species range maps, isotope dietary niche breadth is unlikely to match literature data. This is because diet isotopic dimensions largely depend on the inherent isotopic variance of the resources (Shipley and Matich, 2020), and thus may not adequately reflect the diversity of individual food items in the diet (as reported in the literature), but rather the isotopic diversity of used resources.

Lastly, we assessed whether species isotope niche breadths in the breeding range were related to their long-term population trends as reported in the literature. This is of particular importance for long-distance migrants that show rapid population declines (e.g. Heldbjerg and Fox, 2008; Evans et al., 2012). If species with narrow isotope niches systematically displayed negative population trends, it would indicate that conditions in the breeding grounds are key for the long-term persistence of species.

## Methods

### Feather sampling

Juvenile wing and tail feathers (each grown in the nest; Jenni and Winkler, 2011) from 807 migratory birds belonging to 48 passerine species were sampled at the ‘Bocca di Caset’ and ‘Passo del Brocon’ ringing stations (Province of Trento, Italy, 10°41’ E, 45°51’ N and 11°68’ E, 46°11’ N, respectively) in the central Italian Alps during autumn migration (from late August to the end of October) from 2010 to 2013.

Migratory species pass through the Italian Alps at different times during autumn migration (Pedrini et al., 2008). Three migratory groups were considered: longdistance trans-Saharan migrants (TSM), which are fully migratory and leave their breeding ranges to winter in sub-Saharan Africa; short-distance intra-Palaearctic migrants (IPM), which can be fully or partially migratory and which have their breeding and wintering ranges within the Western Palaearctic; irruptive intra-Palaearctic migrants (IRM), which are partial migrants and residents, but show marked invasive and nomadic movements during non-breeding seasons. Date of capture was also recorded for each individual.

### Stable isotope analysis

Values of δ^2^H, δ^18^O, δ^13^C, δ^15^N were measured for each individual feather. It was possible to determine all four isotope ratios in the majority of individuals. Feathers were first washed in a solvent mixture (diethyl ether-methanol 2:1) prior to analysis (Bontempo et al., 2014). Simultaneous determination of δ^13^C and δ^15^N (i.e. in the same run) was accomplished using a Vario Isotope Cube isotope ratio mass spectrometer (Elementar, Germany). The δ^2^H and δ^18^O values were determined through pyrolysis combustion using a TC/EA (Thermo Finnigan, Bremen, Germany) interfaced with a Delta Plus XP (Thermo Finnigan, Bremen, Germany) continuous-flow isotope-ratio mass spectrometer. Pyrolysis was carried out at 1450°C in a glassy carbon column. The helium carrier flow was 110 ml/min, the GC column was a 1.2 m long molecular sieve 5A, at 110°C. Isotope ratios were expressed in δ-notation against V-PDB (Vienna-Pee Dee Belemnite) for δ^13^C, Air for δ^15^N, V-SMOW (Vienna-Standard Mean Ocean Water) for δ^2^H and δ^18^O. Isotopic values of δ^2^H were calculated using the comparative equilibration approach (Wassenaar and Hobson, 2003, 2006). based on the keratin standards CBS (δ^2^H −197 ‰, δ^18^O = +3.8 ‰) and KHS (−54.1 ‰, δ^18^O = +20.3 ‰). Values of δ^13^C and δ^15^N were calculated against working in-house standards (casein and wheat), which were themselves calibrated against international reference materials using multi-point normalisation: fuel oil NBS-22 (IAEA International Atomic Energy Agency, Vienna, Austria; −30.03 ‰) and sugar IAEA-CH-6 (−10.45 ‰) for δ^13^C, L-glutamic acid USGS 40 (−26.39 ‰ and −4.5 ‰ for δ^13^C and δ^15^N), hair USGS 42 (δ^15^N = +8.05 ‰ and δ^13^C = −21.09 ‰) and USGS 43 (δ^15^N = +8.44 ‰ and δ^13^C = −21.28 ‰) for ^13^C/^12^C and ^15^N/^14^N. Values of δ^13^C were expressed versus V-PDB on a scale normalised against the two reference materials LSVEC (−46.6 ‰) and NBS 19 (+1.95 ‰) (Brand et al., 2014). Values of δ^15^N were expressed versus Air-N2 on a scale normalised using the two reference materials IAEA-N-1 and USGS32, with consensus values of +0.4 ‰ and +180 ‰ (Brand et al., 2014). Method uncertainty (calculated as one standard deviation) was 0.1 ‰ for δ^13^C, 0.2 ‰ for δ^15^N, 0.3 ‰ for δ^18^O and 2 ‰ for δ^2^H.

### Data analysis

We gathered information on migratory behaviour and distance for each bird species from the literature (Del Hoyo et al., 2013). Migration distance was calculated as the distance between the centroids of the breeding and wintering areas. Specifically, distance was estimated as the mean of the two latitudes during breeding minus the mean of the two latitudes during winter (Møller et al., 2008).

To compare empirical estimates of dietary and origin isotopic niche breadth with literature data, we used recently published measures of diet specialisation and climate niche breadth that were derived from literature data and breeding range maps of European birds (Reif et al. 2016). Diet specialisation was expressed as the coefficient of variation in the presence/absence of eight food types in the diet of each species (foliage, fruit, grain, insects, other invertebrates, terrestrial vertebrates, water vertebrates, carrion). Climate niche breadth was measured by overlying the breeding range map of each species to climatic data and calculating the covered temperature range (see Reif et al. 2016 for details).

Information on long-term population trends for each species at the European scale was gathered from the PanEuropean Common Bird Monitoring Scheme (PECBMS) (https://pecbms.info/trends-and-indicators/species-trends/).

Dietary isotopic niche breadth (NB) was expressed as the Bayesian Standard Ellipse Area corrected for small sample size (SEAc) using the isotopic space defined by *δ*^13^C-*δ*^15^N. Similarly, spatially-explicit origin NB was expressed as SEAc from the *δ*^2^H-*δ*^18^O space. The SIBER R-package was used for these calculations (Jackson et al., 2011; Rader et al., 2017).

Differences in NB among migratory groups were examined using generalised least squares (GLS) allowing the variance to differ among groups (with *`varIdent`* function in R). Seasonal trends in *δ*^2^H, *δ*^13^C and *δ*^15^N within each migratory group were assessed using linear mixed-effect models including species as random factor to account for different isotope values and trends across species. We used the *`lme`* function in R allowing both intercept and slope to differ among species. To examine if origin NB was related to migration distance, we used GLS to regress the mean NB values for each species (across all measured individuals) against the estimated migration distance.

Although migration timing can vary within species depending on age and sex, we were not able to consistently record this information from our samples to model it statistically.

## Results

Over the four years of sampling, more than 800 individuals of 48 passerine species were captured. After excluding species with less than 15 individuals, we included isotope data from 715 individuals of 21 species (Tab. S1). These belonged to the trans-Saharan (7 species; 241 individuals), intra-Palearctic (11 species; 335 individuals) and irruptive (3 species; 91 individuals) migratory systems. Overall distribution of species over isotope space is shown in Fig. 1 and 2. Migratory phenology of each species during the autumn passage is shown in Fig.S1 (Supplementary Material), where the temporal patterns in the number of species captured is also shown.

**Fig. 1 -.**
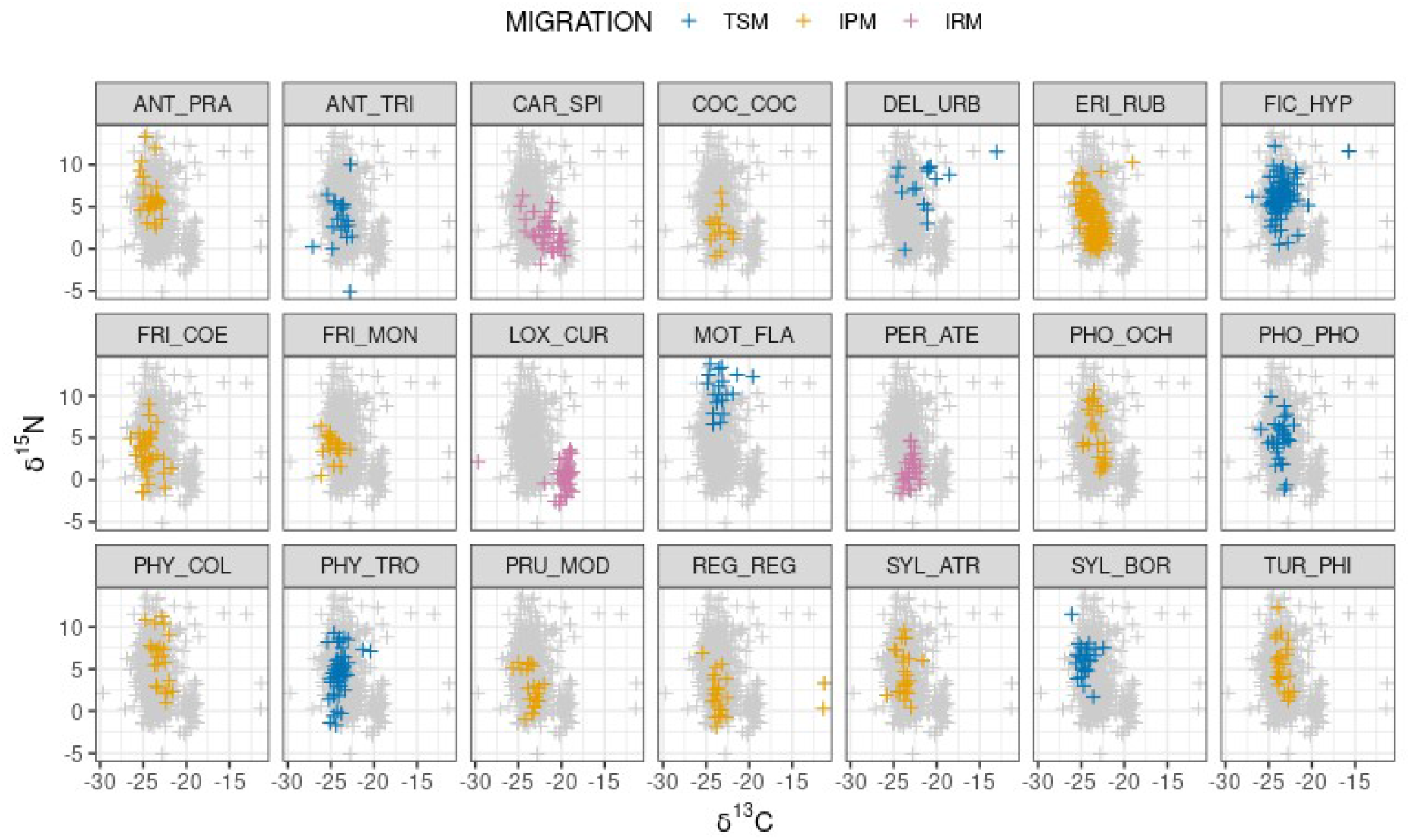
Distribution of each species and individuals over the *δ*^13^C-*δ*^15^N isotope space defining the dietary niche. Grey crosses in the background show the overall space occupied by the species (acronyms provided in Tab.S1). TSM=Trans-Saharan migrants; IPM=Intra-Palearctic migrants; IRM=Irruptive migrants

**Fig. 2 -.**
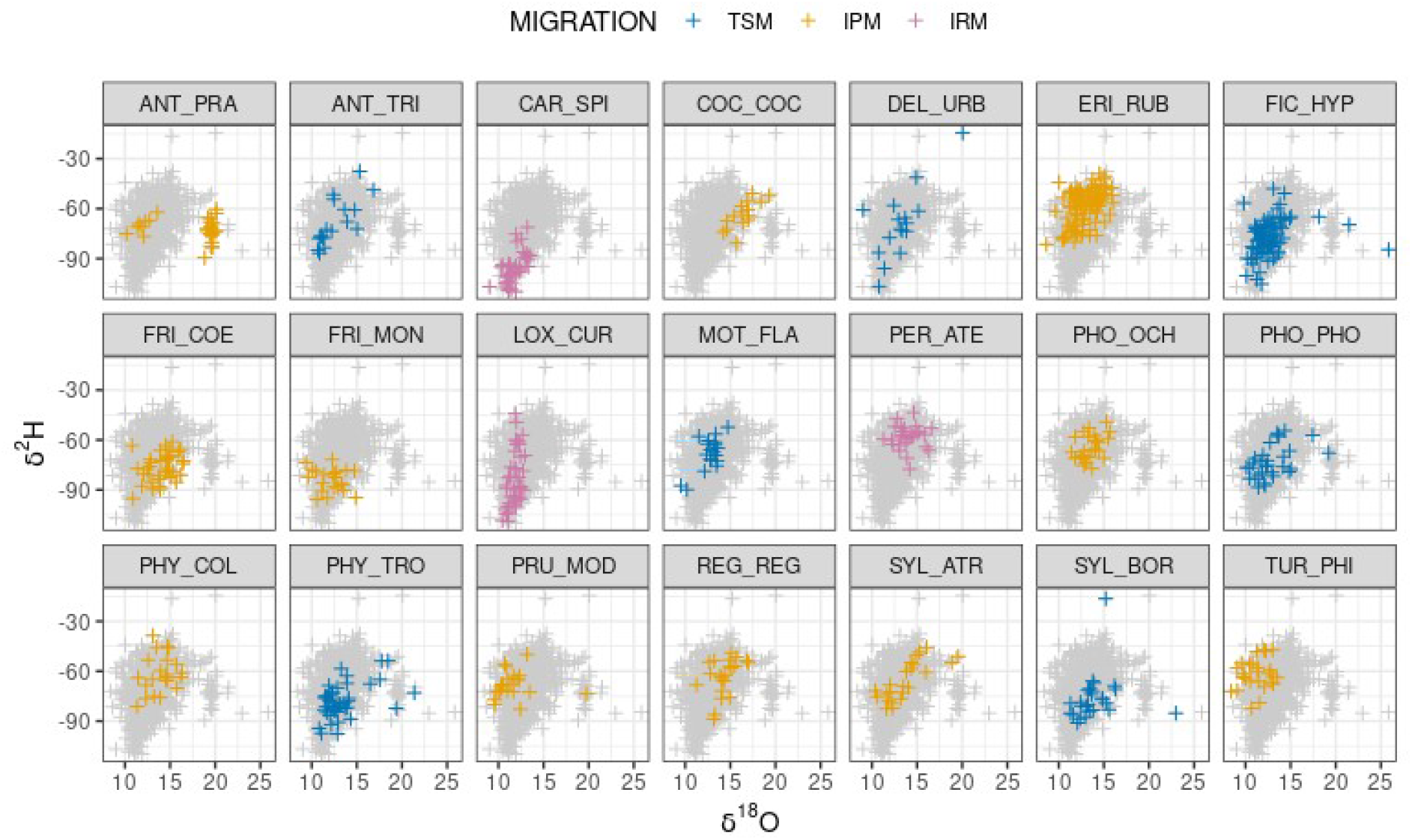
Distribution of each species and individuals over the *δ*^2^H-*δ*^18^O isotope space defining the spatial origin niche. Acronyms as in Fig.1

Seasonal changes in *δ*^2^H (reflecting species breeding latitude) supported the hypothesis that both TSM and IPM groups display a Type-I (chain) migration, whereby northern populations (with lower *δ*^2^H values) reached the Alpine passage later in the season (Fig.3). This pattern appeared consistent among species in each group, with a significant effect of capture day in the *lme* model (p<0.01 for both groups). Mean *δ*^2^H values were evidently lower for TSM, indicating that species in this migratory group tend to breed at higher latitudes than IPM. No seasonal trend in *δ*^2^H was observed in IRM species.

**Fig. 3 -.**
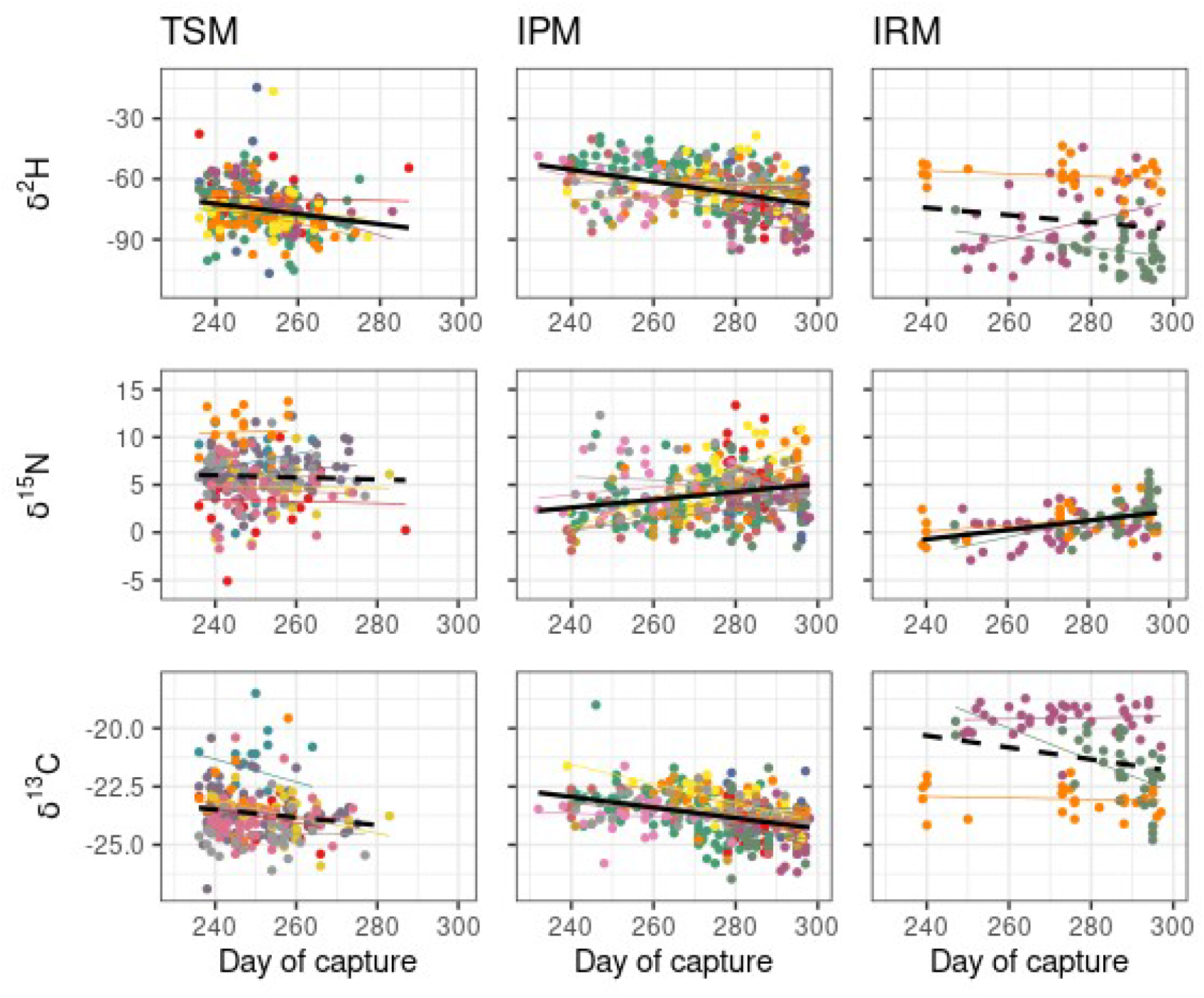
Seasonal trends in *δ*^2^H, *δ*^15^N and *δ*^13^C values for the TSM, IPM and IRM groups. Black fit line indicates overall trend (dashed line=*NS*), while individual species trends are shown in different colours.

A seasonal trend in *δ*^15^N was evident for IPM and IRM species with a positive effect of capture day (*lme* p<0.01). This trend was not observed in TSM, which displayed slightly higher mean *δ*^15^N values (Fig.3). Similarly, a significant negative trend in *δ*^13^C was observed in IPM (p<0.01), which was not significant in either TSM and IRM species. Fig.3 also shows that most TSM species reached the capture site by day 280, whereas the season appeared longer for IPM and IRM. The difference in phenology is also clear from Fig.S1.

Average species dietary isotopic NB did not differ significantly among migratory groups (Fig.4). Conversely, the average isotopic origin niche was significantly broader for TSM, about 30% broader than IPM (GLS model; p=0.03). However, there was relatively large variation in origin NB within TSM. IRM displayed a much narrower origin NB (Fig.4). Excluding the apparent outliers from the analysis of dietary NB (*δ*^13^C-*δ*^15^N; *Regulus regulus* and *Delichon urbicum*), provided qualitatively similar results.

**Fig. 4 -.**
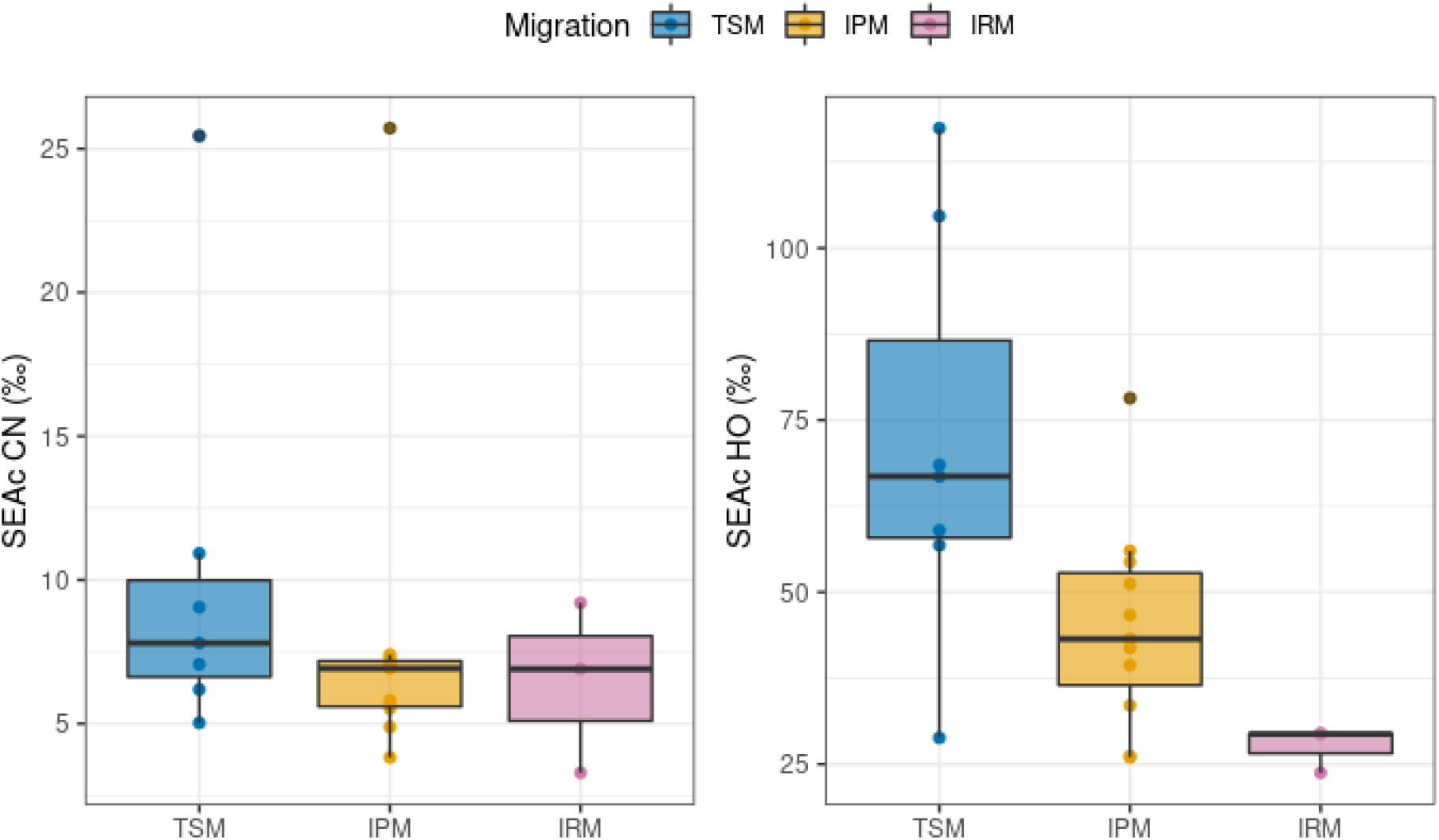
Comparison of the average species dietary and origin isotopic niche breadth among migratory groups.

In addition, migration distance was related to the breadth of species origin niche, although there was considerable variation at larger distances (Fig. 5; GLS; p=0.01). No relationship was observed between dietary isotopic NB and migration distance (Fig.S2).

**Fig. 5 -.**
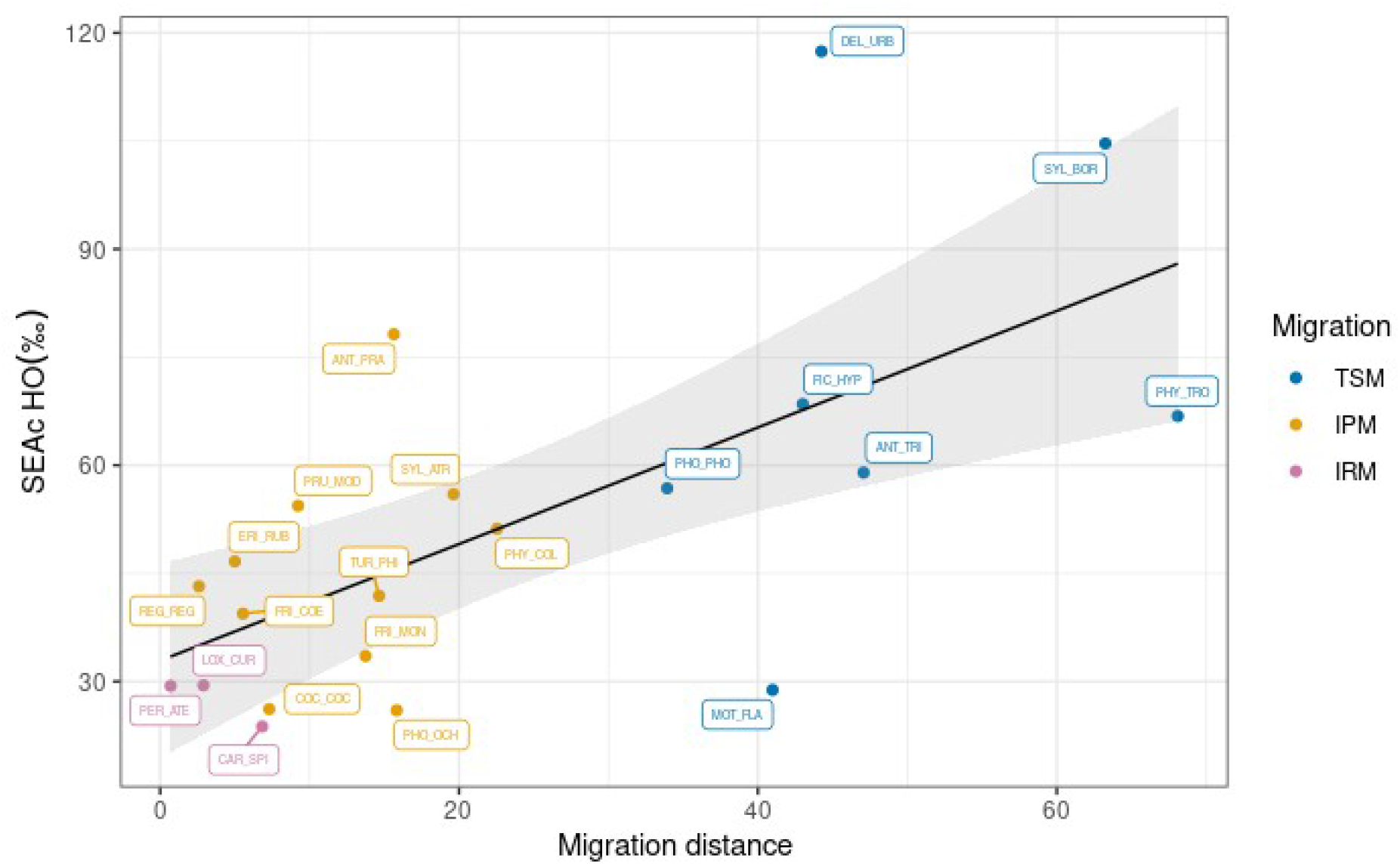
Relationship between species origin niche breadth and migration distance, defined as the latitudinal distance between the centroid of the breeding and wintering ranges. Species identity is labelled.

We compared our estimates of species dietary and origin isotopic niche with those recently derived from species distributional data. We found that origin NB well matched measures from range maps (Fig. 6), with a positive linear correlation (p=0.02). In addition, climate niche breadth derived from range maps well reflected the upper distribution of the empirical isotope origin niche. This is evident from the significant quantile regression at q=0.8 (p<0.01). Conversely, the dietary isotopic NB was not related to diet specialisation, as derived from the literature.

**Fig. 6 -.**
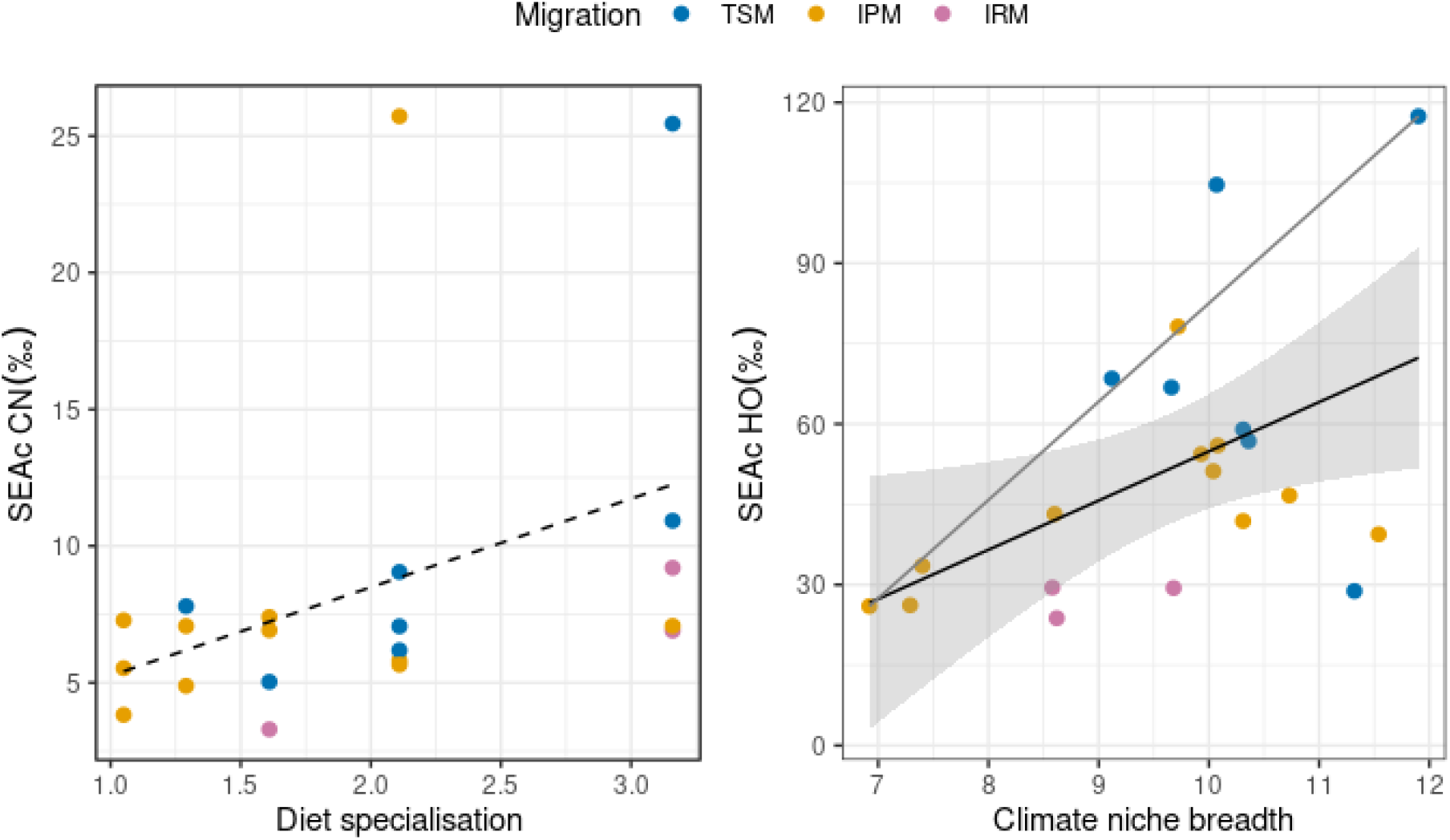
Relation between empirical dietary and origin isotopic niche breadth with literature-derived niche parameters (from Reif et al 2016). Dashed line = *NS* relation; tick black line = significant regression fit; thin grey line = significant quantile regression fit at q=0.8

Finally, we appraised whether long-term population trends were related to the empirical isotope niches. Neither dietary not origin isotopic niche breadth were related to species population trends as reported in the PECBMS portal (Fig.S3).

## Discussion

We used juvenile feathers of multiple bird species intercepted along an autumn migratory flyway as ‘remote carriers’ of isotope niche information from Western Palearctic breeding areas. This allowed quantifying niche parameters from species that evolved different migratory strategies. In addition, despite potential confounding factors related to variance in local isotopic baselines, including individuals from different populations and communities allowed the estimation of species niches that were less influenced by local biotic interactions (e.g. competition), and likely more representative of the realised niche dimensions of the species.

Our results generally supported our initial hypotheses, but also revealed a poor association between isotope niche dimensions and literature-derived niche parameters and long-term population trends.

Seasonal trends in feather *δ*^2^H indicated that individuals breeding at higher latitudes (displaying lower *δ*^2^H values) reached the sampling locations later in the season, with patterns that appeared consistent among species and across the two major migratory groups (TSM, IPM). This type of migration, often called Type I (as opposed to Type-II, whereby northern populations leap-frog southern populations during migration), has also been observed across a range of Nearctic-Neotropical migrants (Dunn et al. 2006, Cardenas-Ortiz et al., 2020), and appears common in Afro-Palearctic migrants as well (Briedis et al., 2016, 2020). Different migration timing of conspecific populations contributes to limit competition among individuals and likely occurs at the species level too. Assessing temporal segregation in the use of migratory stopover helps identify the species most likely to co-occur and compete locally for resources. In our study site, a few species showed complete segregation (Fig.S1), but high overlap was otherwise evident with up to nine migratory species captured in a single day. Bottlenecks can indeed concentrate large numbers of individuals and species during a short period, which not only heightens intra- and inter-specific competition, but also increases vulnerability to localised events and disturbance (Cardenas-Ortiz et al., 2020). The identification and conservation of migratory stopover and bottlenecks are therefore key for the persistence of many declining populations (Heiss, 2013; Buechley et al., 2018; Cardenas-Ortiz et al., 2020).

Seasonal trends observed in both *δ*^15^N and *δ*^13^C in IPM is indicative of possible shifts in the availability or quality of resources for these species. However, these patterns could be explained by different non-mutually exclusive mechanisms. For instance, seasonally rising values of *δ*^15^N could be associated with the increased availability of adult insects (as opposed to larvae or other arthropods with lower *δ*^15^N values) as the longer breeding season of IPM progresses. Conversely, the shorter season of TSM and their strictly insectivorous diet would result in more seasonally stable *δ*^15^N values. Alternatively, a geographic explanation is also plausible. The lower latitude breeding range of IPM, as well as IRM (compared to TSM), implies that early captures were dominated by individuals from the Alpine/central-Europe/Carpathian areas at high elevations. This could be associated with higher *δ*^13^C (Hobson et al., 2004) and possibly with lower *δ*^15^N due to lower precipitation compared to northern Europe (Cortesi et al., 2012). It is also possible that the extent of anthropogenic nitrogen input from agriculture is lower in mountainous regions, which could contribute to lower *δ*^15^N in early captures. It is known, in fact, that anthropogenic sources such as fertilisers can alter and mostly increase nitrogen isotopic baseline values (Shipley and Matich, 2020).

We found that species breeding origin niche was, on average, broader in long-distance trans-Saharan compared to intra-Palearctic migrants. Long-distance migration is generally thought to have evolved from ancestor species with broad habitat niches: individuals able to exploit a wider range of conditions during migration or at wintering grounds should be selectively advantaged over more specialised ones (Levey and Stiles, 1992; Cresswell, 2014; Reif et al., 2016). Empirical support for this hypothesis is limited (Salewski et al., 2003; Jones et al., 2010; Laube et al., 2015), and our data thus provide some additional evidence. However, the link between niche breadth and migration patterns remains unclear, with some evidence suggesting the opposite view that narrower habitat or diet niche may have favoured long-distance migrations (e.g. Brändle et al., 2002), because specialists are forced to leave an area when conditions become unfavourable or resources scarce. Yet, our data do not show any correlation between migration distance and dietary niche breadth, providing no support for this alternative hypothesis. It has to be kept in mind, however, that our estimates of ecological niches reflect the breeding range of species (i.e. isotopes from juvenile feathers grown at nest), and the inclusion of information from wintering grounds would have provided a more complete and likely different picture. This would require measuring isotope ratios in body tissues with different turnover or growth rates (Hahn et al., 2013), or collecting feathers grown in winter (Steenweg et al., 2017). However, for most passerine species, information on wintering locations is still limited, especially for the Afro-Palearctic migration system (Finch et al., 2017; Somveille et al., 2019). Similarly, the extent to which migratory species track their climate and habitat niche between breeding and wintering grounds is poorly understood and likely to vary among species (Somveille et al., 2019; Ponti et al., 2020). Most assessments of seasonal niche shifts during migration relied on large-scale climatic and land-use data (Gómez et al., 2016; Zurell et al., 2018; Ponti et al., 2020), while those employing tissues isotopes are rare and mostly focussed on a few key species to appraise migratory connectivity (Rubenstein, 2002; Yohannes et al., 2007; Hahn et al., 2013). Therefore, it is difficult to evaluate whether and to what extent our estimates of isotope niche breadth would have differed had we included wintering isotope signals.

We used recently published data from Reif et al. (2016) to examine the extent to which isotope niches from our sampled populations reflected diet specialisation and climate niche breadth reported at the species level. As expected, dietary isotopic niche did not match the breadth or diversity of consumed food items reported in the literature. This further supports the notion that the breadth of dietary isotope niche reflects the isotopic variance of the resources and should not be considered synonymous of species diet (Shipley & Matich, 2020). Conversely, estimates of spatially-explicit breeding origin niche correlated significantly, albeit weakly, with literature data. In particular, measures of climatic niche from range maps appeared to better predict the upper distribution (quantile: 0.8) of our isotope niche, rather than its central response. A possible interpretation is that niche estimates derived from range maps necessarily reflect the maximum expected value for a given population. In other words, if we assume that a correlation exists between spatially-explicit origin niche inferred from isotope and range maps, any sampled populations should display isotope niches that are either equal or narrower than those derived from species full range maps. This provides additional evidence that species bi-dimensional *δ*^2^H-*δ*^18^O isotope space is a good proxy of their geographic and climatic niche dimensions (e.g. Rader et al., 2017).

Among the key motivations for appraising niche requirements of migratory species, is that their populations appear to be declining more rapidly than residents (Sanderson et al., 2006; Heldbjerg and Fox, 2008; Evans et al., 2012; Ockendon et al., 2012). The mechanisms underpinning these trends are still unclear and we examined whether narrow isotope breeding niches were associated with stronger population declines. We found no correlation between either dietary or origin breeding niche and long-term population trends as reported in the Pan European Common Bird Monitoring Scheme. This is not surprising as there is increasing evidence that environmental conditions and dynamics in non-breeding areas are key for the long-term persistence of populations (Keller and Yahner, 2006; Sanderson et al., 2006; Morrison et al., 2013). In particular, habitat change and degradation in tropical wintering areas appear to be a major cause of decline in migrant species (Bennett et al., 2018; Şekercioğlu et al., 2019). This further emphasises that quantifying both breeding and wintering niche parameters would provide insight not only on basic ecological questions such as the degree of niche tracking, but also on conservation requirements of long distance migrants (Hahn et al., 2013).

We used multiple stable isotopes to quantify and compare ecological niche parameters between short- and long-distance migrants. We showed that autumn migratory bottlenecks, such “Bocca di Caset” and “Passo del Brocon” in the Italian Alps, not only represent important stopovers for many migrants of the Western flyway (Briedis et al., 2020), but also strategic natural laboratories to examine migratory patterns across multiple species.

Our results illustrate that long-distance trans-Saharan migrants, mostly breeding at higher latitudes than intra-Palearctic migrants, reach the migratory stop-over sooner. This is in line with the notion that early departure allows trans-Saharan migrants to reach the Sahel zone at the peak of vegetation greenness, when feeding conditions are optimal (Newton, 2007; Thorup et al., 2017). Moreover, in both migratory groups, the onset of breeding for northern populations appeared delayed, likely due to climatic constraints, so that the timing of migration was proportionally shifted later in the season. The delayed timing of northern populations can apparently carry over to the entire annual cycle (Briedis et al., 2016), and likely contributes to limit intraspecific competition. This type of migration pattern appears fairly common in many Neartic-Neotropical migrants too (Dunn et al. 2006; Cardenas-Ortiz et al., 2020). Our data also provide some support to the hypothesis that broad ecological niches are linked to long-distance migration, although this was only evident for the breeding origin niche dimensions, and requires further testing.

The use of multiple isotope ratios in animal tissues has allowed the quantification of both dietary and spatially-explicit niche aspects (Shipley and Matich, 2020). However, the relation between the isotopic ecological niche and other descriptors of species niche (e.g. based on feeding habits, distribution patterns), or conservation status remains vague (Rader et al., 2017; Shipley and Matich, 2020). Here, we found that origin isotopic niche reflected the climatic range extent of the species, illustrating how δ^2^H and δ^18^O measurements can provide insight into origins related to bio-climatic and geographic isotopic patterns. However, the breadth of isotope niches was not able to explain the long-term population trends of the species at the European scale. Appraisal of the complete multi-seasonal niche dimension of longdistance migrants is therefore needed to link ecological information to conservation actions for these declining species.

## Acknowledgements

Funding to A.F. was provided by FEM – Fondazione Edmund Mach, Research and Innovation Centre and MUSE – Science Museum of Trento (Accordo di Programma – Provincia Autonoma di Trento). This study complies with the current Italian laws regulating scientific research on animals. The authors gratefully acknowledge G. Bogliani and S. Tenan, for their suggestions in the early phase of this study, and A. Tonon and L. Ziller for the analytical support, S. Noselli, F. Rizzolli and F. Rossi for field work.

## Notes

**Declarations** Funding. Funding to A.F. was provided by FEM – Fondazione Edmund Mach, Research and Innovation Centre and MUSE – Science Museum of Trento (Accordo di Programma – Provincia Autonoma di Trento).

The Authors declare no conflict of interest.

### Competing Interest Statement

The authors have declared no competing interest.

