## Supplementary documents for "Multidimensional natal isotopic niches reflect migratory patterns in birds"

### Supplementary Material

Table S1 – List of bird species included in the analyses. Latin and English names are given, with acronyms used in the plots.

| LATIN | ENGLISH | ACRONYM |
|---|---|---|
| *Anthus pratensis* | Meadow Pipit | ANT_PRA |
| *Anthus trivialis* | Tree Pipit | ANT_TRI |
| *Coccothraustes coccothraustes* | Hawfinch | COC_COC |
| *Delichon urbicum* | Northern House Martin | DEL_URB |
| *Erithacus rubecula* | European Robin | ERI_RUB |
| *Ficedula hypoleuca* | Pied Flycatcher | FIC_HYP |
| *Fringilla coelebs* | Common Chaffinch | FRI_COE |
| *Fringilla montifringilla* | Brambling | FRI_MON |
| *Loxia curvirostra* | Red Crossbill | LOX_CUR |
| *Motacilla flava* | Western Yellow Wagtail | MOT_FLA |
| *Periparus ater* | Coal Tit | PER_ATE |
| *Phoenicurus ochruros* | Black Redstart | PHO_OCH |
| *Phoenicurus phoenicurus* | Common Redstart | PHO_PHO |
| *Phylloscopus collybita* | Common Chiffchaff | PHY_COL |
| *Phylloscopus trochilus* | Willow Warbler | PHY_TRO |
| *Prunella modularis* | Dunnock | PRU_MOD |
| *Regulus regulus* | Goldcrest | REG_REG |
| *Spinus spinus* | Eurasian Siskin | CAR_SPI |
| *Sylvia atricapilla* | Eurasian Blackcap | SYL_ATR |
| *Sylvia borin* | Garden Warbler | SYL_BOR |
| *Turdus philomelos* | Song Thrush | TUR_PHI |

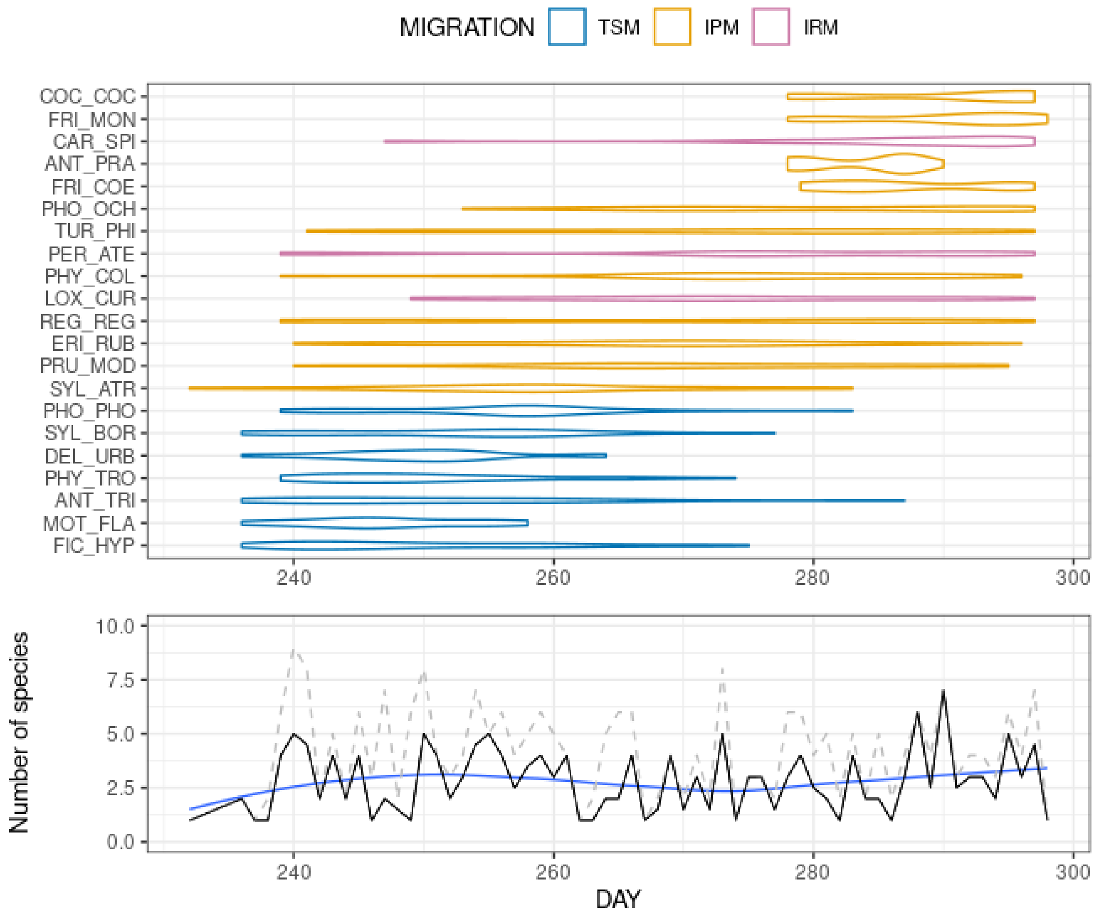


**Fig.S1** - Migratory phenology of the 21 species during the autumn migration over the Italian Alps. Violin plots were built combining four years of sampling (2010-2013).

Lower panel shows the median (across years; black line) and maximum (grey dashed line) number of species captured each day. A LOESS smooth of the median is also shown.

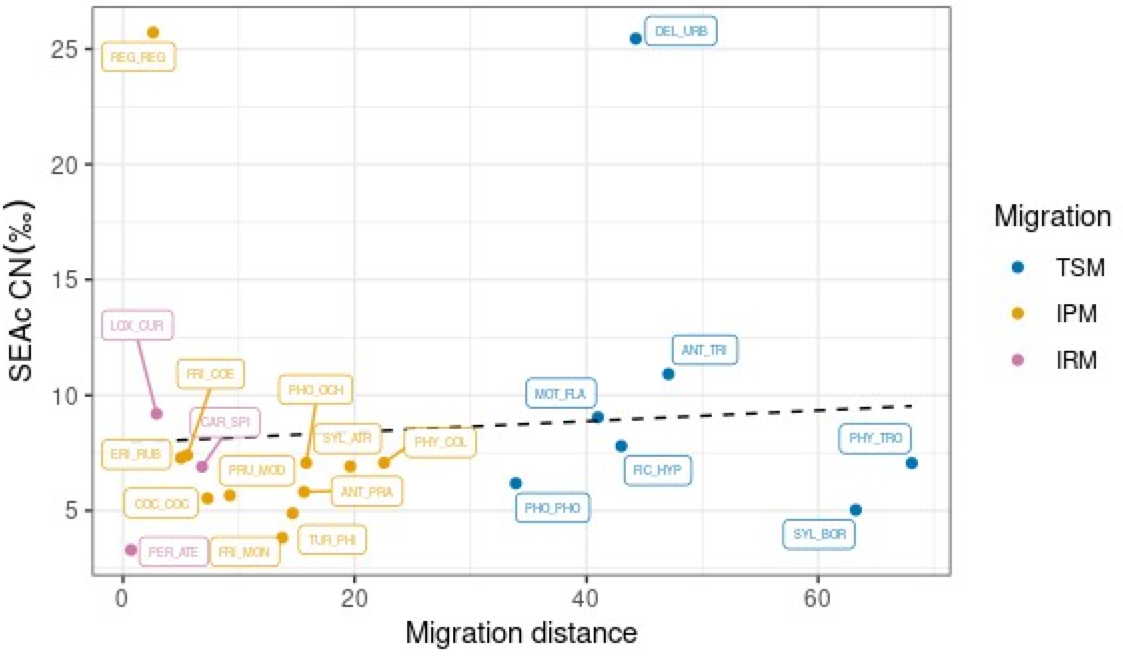


**Fig S2**- Relationship between species dietary NB and migration distance, defined as the latitudinal distance between the centroid of the breeding and wintering ranges. Species are labelled.

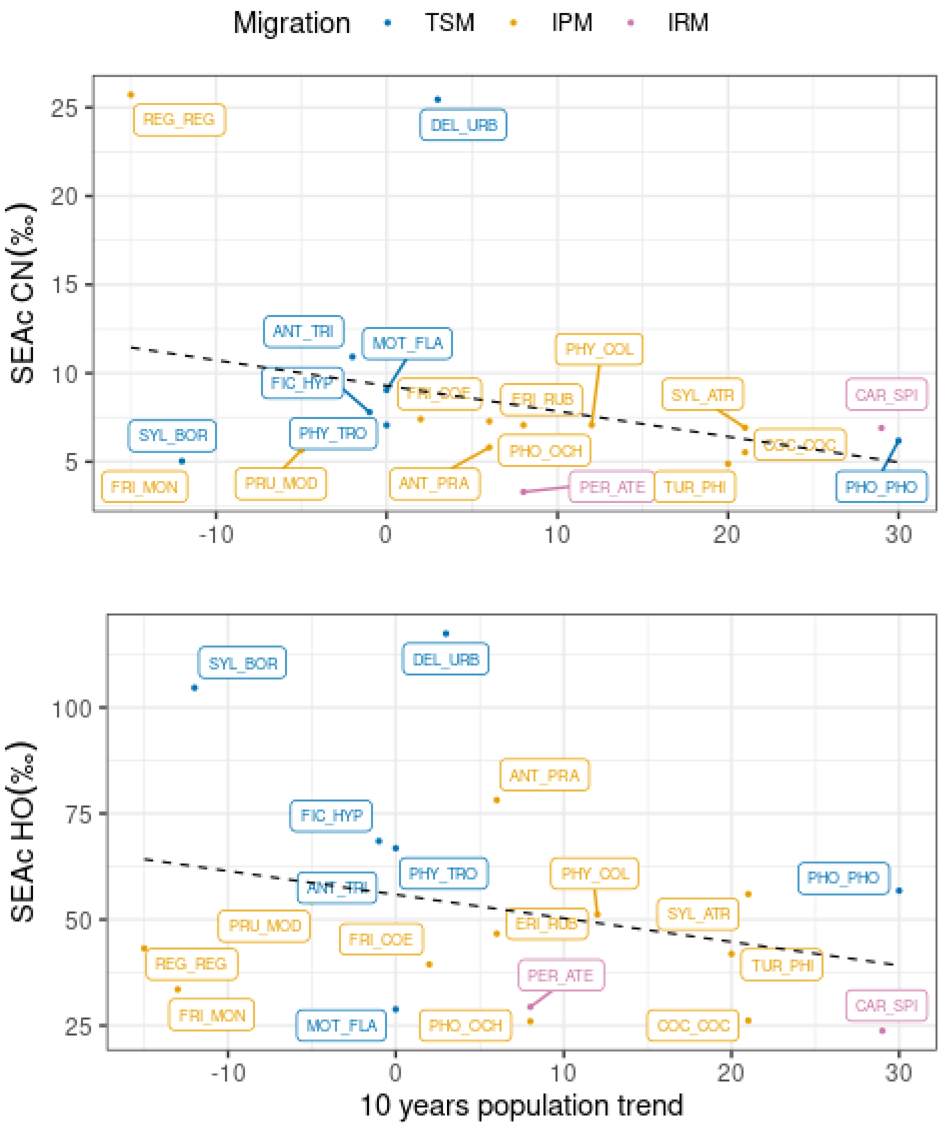


**Fig.S3** - Relationship between isotopic dietary and origin breeding niche breadth and population trends as reported in the Pan European Common Bird Monitoring Scheme. Dashed line indicates non-significant trends. Species are labelled.
